# Dissociable neural effects of temporal expectations due to passage of time and contextual probability

**DOI:** 10.1101/769372

**Authors:** Ana Todorovic, Ryszard Auksztulewicz

## Abstract

The human brain is equipped with complex mechanisms to track the changing probability of events in time. While the passage of time itself usually leads to a mounting expectation, context can provide additional information about when events are likely to happen. In this study we dissociate these two sources of temporal expectation in terms of their neural correlates and underlying brain connectivity patterns. We analysed magnetoencephalographic (MEG) data acquired from N=24 healthy participants listening to auditory stimuli. These stimuli could be presented at different temporal intervals but occurred most often at intermediate intervals, forming a contextual probability distribution. Evoked MEG response amplitude was sensitive to both passage of time and contextual probability, albeit at different latencies: the effects of passage of time were observed earlier than the effects of context. The underlying sources of MEG activity were also different across the two types of temporal prediction: the effects of passage of time were localised to early auditory regions and superior temporal gyri, while context was additionally linked to activity in inferior parietal cortices. Finally, these differences were modelled using biophysical (dynamic causal) modelling: passage of time was explained in terms of widespread gain modulation and decreased prediction error signalling at lower levels of the hierarchy, while contextual expectation led to more localised gain modulation and decreased prediction error signalling at higher levels of the hierarchy. These results present a comprehensive account of how independent sources of temporal prediction may be differentially expressed in cortical circuits.

**HIGHLIGHTS:**

- Predictability of tone onset times affects auditory network connectivity
- Foreperiod and distribution of events in time have dissociable neural substrates
- Decreased prediction error at different levels of cortical hierarchy

## INTRODUCTION

If we know that we will hear a tone between this instant and a fixed future moment, but not exactly when, then, as we wait for the tone to arrive, its probability of occurring at the next instant will grow. The possible onset times of the tone might also have a particular temporal distribution, for example, tones might be more frequent within a narrow time window. The probability of such a tone occurring at a given moment will be a combination of the distribution of onset times and the probability of tone onset increasing over time (Nobre, Correa & Coull, 2007; Janssen & Shadlen, 2005). This combined computation is known as the hazard rate.

The brain tracks temporal hazard rates using a wide network of areas. Mounting temporal expectation correlates with BOLD activity in early sensory cortex, the cerebellum, as well as fronto-parietal regions including the inferior parietal cortex (IPC), supplementary motor area (SMA), and frontal areas (Bueti, et al., 2010). Electrophysiological studies have also identified that early sensory regions (Ghose & Maunsell, 2002), the SMA (Akkal, et al., 2004), and lateral intraparietal cortex (Janssen & Shadlen, 2005; Leon & Shadlen, 2003) are all sensitive to temporal expectation.

While temporal expectation modulates neuronal activity over multiple regions including sensory and frontal cortices, other areas, such as the superior temporal gyrus (STG), SMA, and IPC may be more sensitive to the final integration of passage of time and contextual information (Muller & Nobre, 2014). For example, one study found that activity only in prefrontal, but not in sensory regions, could be linked with faster reaction times in the presence of temporal expectation (Vallesi, et al., 2009). A different study found that while the superior temporal gyri (STG), SMA, and prefrontal regions were all involved in time estimation, only the STG was involved in estimating the duration of one stimulus in comparison to another stimulus (Coull, et al., 2008), showing additional sensitivity to context. A similar dissociation has been found more recently within the frontoparietal network, where the IPC showed sensitivity to fixed contextual temporal probabilities while activity in the intraparietal sulcus and inferior frontal regions was modulated by hazard rate (Coull, et al., 2016).

Apart from estimating which brain areas are involved in tracking hazard rates, an important question is how they communicate in the process. A large body of literature finds that when stimulus events are predictable, less neural activity is needed to process them (Kok, Jehee, and de Lange 2012; Wacongne, et al. 2011; Todorovic, et al. 2011), and this includes predictability of stimulus occurrence in time (Alink, et al. 2010; Costa-Faidella, et al. 2011; Fischer, Plessow, and Ruge 2013; Lange 2013). Predictive coding models posit that this decrease in neural activity rests on recurrent message passing between areas along the cortical hierarchy (Friston 2005; Lee and Mumford 2003; Spratling 2010), where less predictable stimuli would correspond to a network with more feedforward activation (corresponding to prediction errors being sent forward), less feedback inhibition (corresponding to predictions being sent back) and less intrinsic inhibition (corresponding to the gain or precision of prediction errors).

In this study we asked whether the passage of time and the distribution of events in time are jointly tracked in the brain as a unitary probabilistic computation, or whether, alternatively, they correspond to two probabilistic computations with different correlates in the auditory network. We additionally looked at how the brain areas involved in these two elements of tracking the hazard rate mutually interact. We found that temporal expectation of tones could be localised to sources corresponding to the primary auditory cortex (A1), superior temporal gyrus (STG) and inferior parietal cortex (IPC). The hypothesis of two separate probabilistic computations better described our data, with passage of time having an earlier effect than temporal distribution of events, as well as affecting connectivity in a broader network, that partially diverged from predictive coding models.

## MATERIALS AND METHODS

### 2.1 Participants

Twenty-five healthy participants (17 female, mean ± SD age 23.7 ± 7.8 years) enrolled in the study upon written informed consent, and one was later excluded due to excessive measurement noise. All experimental procedures were approved by the local ethics committee (Committee on Research Involving Human Subjects, Region Arnhem-Nijmegen, The Netherlands) and were conducted in accordance with the Declaration of Helsinki. Participants had normal hearing, normal or corrected-to-normal vision, and no history of neurological or psychiatric disorders. Data from one participant were excluded from analysis due to excessive artefacts in MEG recordings (>30% trials).

### 2.2 Experimental paradigm

Stimuli consisted of brief pure tones (5 ms long, frequency 1000 Hz or 1200 Hz, ~70 dB SPL) presented binaurally via MEG-compatible air tubes. Stimulus delivery was controlled using Presentation software (Neurobehavioral Systems).

In each trial (see Figure 1A for an outline of the experimental paradigm; for more details see: Todorovic, et al., 2015), a central fixation cross was presented for 2-4 s, followed by a standard tone (1000 Hz) presented twice, with one of five inter-stimulus intervals (ISI) between the two tones (250, 375, 500, 625, or 750 ms). After the offset of the second tone, the fixation cross remained on the screen for 0.5-1 s and was subsequently replaced by a blank screen for 1.5-2 s. Occasionally, standard tones were replaced by deviant tones (1200 Hz; 9% trials) and participants were instructed to press a button with the right index finger as soon as they heard a deviant tone.

**Figure 1.**
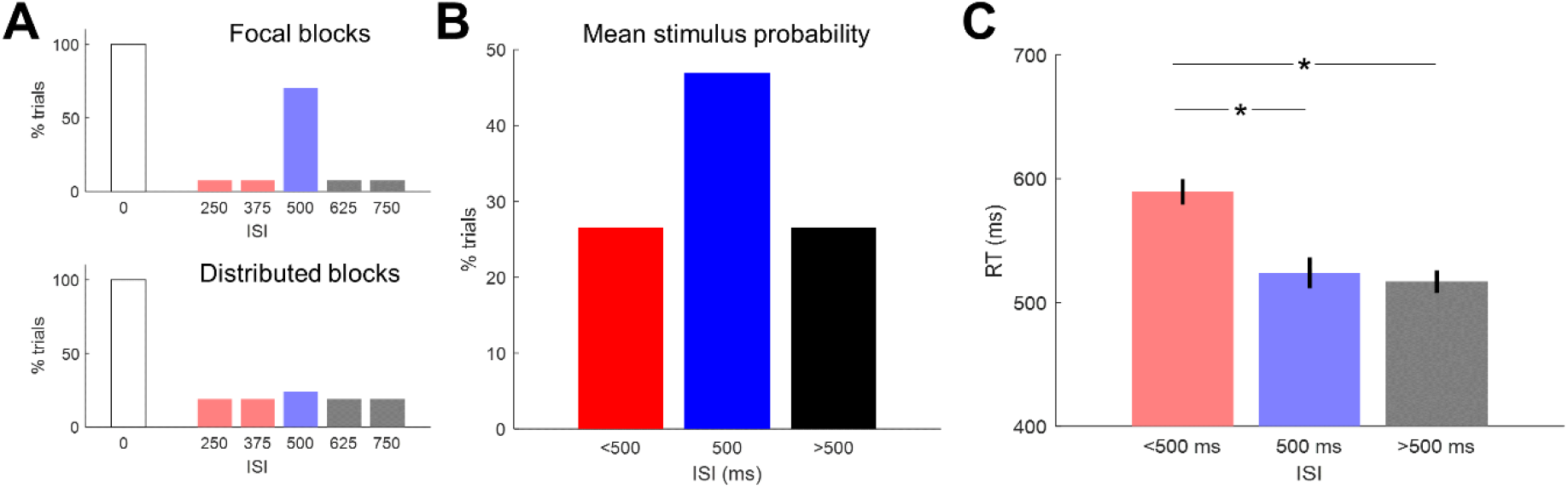
Experimental stimuli and behavioural results. ***(A)*** Following a first tone marking the beginning of a trial (white bar), another tone was presented at a variable inter-stimulus interval (ISI). In focal (distributed) blocks, the second tone was delivered at ISI = 500 ms with 70% (24%) probability among five possible ISIs. Occasionally, either of the two tones was presented with a deviant pitch; participants were instructed to detect these deviants with a speeded button press. ***(B)*** Averaging across focal and distributed blocks, the probability of delivering the second tone at ISI = 500 ms was higher (47%) relative to shorter or longer ISIs (26.5%). ***(C)*** Effects of ISI on RTs: deviant tones presented at short latencies (<500 ms) were detected slower than deviant tones presented at ISI equal or longer than 500 ms. Error bars denote SEMs across participants.

In addition to the influence of passage of time on temporal expectations (i.e., the longer the foreperiod after the first tone, the more likely the presentation of the second tone), we also varied the width of the distribution of second tone presentation at each ISI. In blocks with *focal* temporal expectation, the second tone was played following an ISI of 500 ms in 70% trials, and following other ISIs in 7.5% trials each. As a result, participants could build up a narrow expectation of the second tone being presented 500 ms after the first tone. Conversely, in blocks with *distributed* temporal expectation, the second tone was played following an ISI of 500 ms in 24% trials, and following other ISIs in 19% trials each. In these blocks, participants could expect the second tone to be presented at any ISI with roughly the same probability. In total, four blocks with distributed temporal expectation (corresponding to 88 tones at the ISI = 500 ms and 252 tones at the remaining ISIs), and two blocks with focal temporal expectation (corresponding to 120 tones at ISI = 500 ms and 52 tones at the remaining ISIs) were administered in each participant. The order of blocks was counterbalanced across subjects. Finally, the experimental paradigm included an orthogonal manipulation of temporal attention, whereby participants were instructed to attend to the first or second tone and press a button following a deviant tone presented in the attended position.

### 2.3 Behavioural analyses

We log-transformed the single-trial deviant detection RTs to normalise RT distribution. Trials with RTs outside of the mean±2*SD range were excluded from analysis. Individual participants’ mean RTs to deviants presented in the second position, calculated separately for ISIs shorter than, equal to, and longer than 500 ms respectively, were entered into a repeated-measures ANOVA with a 3-level factor ISI. For plotting purposes, data were back-transformed by exponentiating the individual mean RTs.

### 2.4 MEG acquisition

Magnetic fields induced by neural activity were measured using a whole-head MEG (VSM/CTF Systems) with 275 axial gradiometers. To enable continuous head localization, coils were placed at the nasion and right and left preauricular points, and monitored continuously during the experiment. Electro-oculogram (EOG) and electrocardiogram (ECG) were recorded to guide eye blink and heart beat artefact rejection, using 10-mm-diameter Ag— AgCl surface electrodes.

### 2.5 Event-related field analysis

Continuous data were downsampled from 1200 Hz to 300 Hz, notch-filtered at 50 Hz and high-pass filtered at 0.1 Hz using a two-pass Butterworth filter. Artefacts induced by eye blinks were removed by subtracting two principal spatiotemporal modes associated with eye blinks (Ille, et al. 2002). Corrected sensor data were low-pass filtered at 48 Hz.

Our aim was to compare second tones, that followed the first tone with varying ISIs, to each other. Given that tones at shorter ISIs arrived while the first tone was still being processed, we needed to assess what the evoked field to the second tone at all ISIs would look like in the absence of the first tone. To this end, rather than epoching the data, we modelled the evoked responses using convolution modelling in continuous sensor data estimated for the entire session using several regressors. Convolution modelling (Figure 2; cf. Litvak, et al., 2013; see Spitzer, et al., 2015; Auksztulewicz, et al., 2017 for applications) is equivalent to testing for the effects of independent stimulus events at each point in peristimulus time but at the same time allows for overlapping responses to successive trials, in the same way one would model fMRI time-series. Our principal aim was to estimate the evoked responses to tones presented at different ISIs relative to the first tone. To increase the number of events in each regressor, we pooled over ISIs shorter than 500 ms and longer than 500 ms respectively, resulting in three groups of ISIs (short, middle, long). Thus, the regressors included experimental regressors coding for tone onsets (standard tone in the first position; deviant tone in the first position; standard tone in the second position at ISI lower than, equal, and higher than 500 ms respectively; deviant tone in the second position pooled over all ISIs) and motor responses (button presses) as well as nuisance regressors. Tone regressors were further convolved with a binary regressor coding for temporal attention (on or off the tone of interest). Nuisance regressors included EOG time-series and its temporal derivative, as well as the ECG time-series, to further reduce artefacts associated with eye movements and heart blinks. Furthermore, based on continuous head position measurement inside the MEG scanner, we calculated 6 movement parameters (3 translations and 3 rotations; cf. Stolk, et al., 2013), which were used as further nuisance regressors.

**Figure 2.**
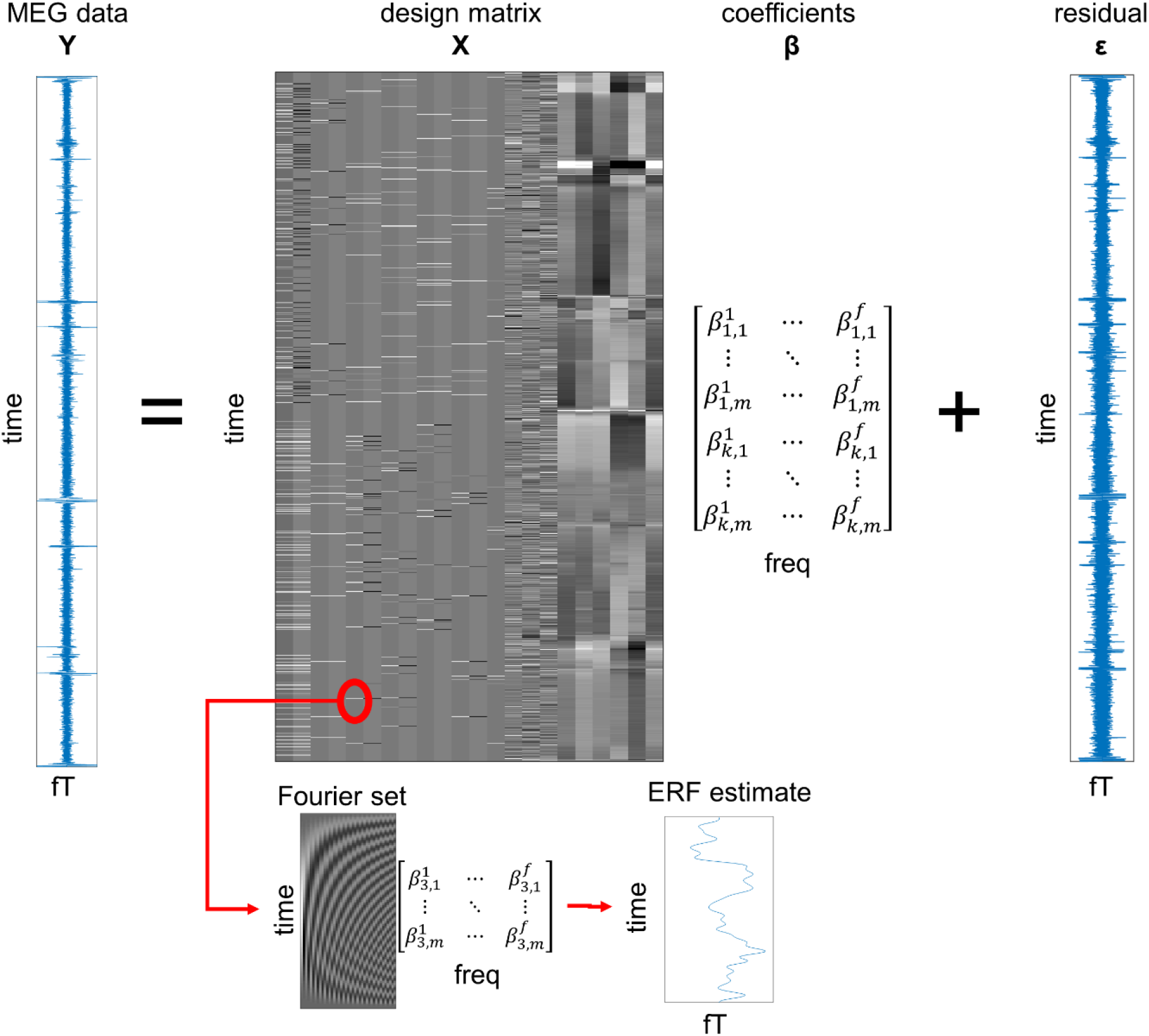
Convolution modelling. Individual participants’ MEG data from the entire experiment (without epoching) were modelled in a general linear model. The design matrix specified the event and nuisance regressors (first 12 columns, left to right: standard onset at ISI = 0 ms, deviant onset at ISI = 0 ms, standard onset at ISI < 500 ms, standard onset at ISI = 500 ms, standard onset at ISI > 500 ms, deviant onset at ISI > 0 ms; each with a corresponding modulation by attention on/off tone; further columns: button press, EOG and its temporal derivative, ECG, 6 motion regressors). Since each regressor was modelled as a Fourier time series (inset below), the ensuing Fourier coefficients can be summarised as a deconvolved ERF response to each event type and/or parametric regressor.

Single participants’ evoked responses were modelled with a time range of 0 ms to 1000 ms relative to events of interest (tone onsets), and other events in the experimental session (button presses). While the actual window of interest encompassed the first 500 ms following tone onsets, we modelled evoked responses up to 1000 ms post-onset in order to deconvolve the influence of neural responses to the first tone from the neural responses to the second tone, especially at short ISIs (shorter or equal to 500 ms). Each event regressor was convolved with a 20-order Fourier basis set with a Hanning window, allowing for an estimation of evoked responses (specifically, their regressor coefficients) per MEG channel with the time-course estimates modulated up to 20 Hz. The resulting spatiotemporal maps of regressor coefficients were converted into 3D images and entered into second-level general linear models (GLMs).

At a group level, we estimated the modulation of the tone-evoked neural responses by two different kinds of temporal expectation: passage of time (monotonically increasing with increasing foreperiod) and tone onset distribution (following an inverted U-shape, approximating the average contextual expectation across the focal and distributed blocks; Figure 1B). The foreperiod effect was thus tested in an F-contrast defined as [1 2 3; −1 −2 −3] over the design matrix columns coding for tones presented in the second position at early, middle, and late ISIs respectively. Conversely, the contextual effect F-contrast was defined as [−1 2 −1; 1 −2 1] over the same columns. Significant effects were inferred under the assumptions of random field theory (Kilner, et al. 2005) after thresholding the 3D (time x 2D topography) statistical parametric maps at *P* < 0.001 (peak-level, uncorrected) and correcting P-values for multiple comparisons based on cluster size at a family-wise error rate at *P* < 0.05.

### 2.6 Source reconstruction

We identified significant effects of foreperiod and context for two separate clusters (250-500 ms, corresponding to the effect of foreperiod; and 425-500 ms, corresponding to the effect of context; see Results 3.2). We then based source reconstruction on a multiple sparse priors algorithm (Friston, et al. 2008) performed on these two time windows. For each time window, the estimated sensor-level evoked responses were first used to calculate contrast time-series, corresponding to the linear approximation to the passage-of-time and the inverted U-shaped approximation to the distribution effect. The resulting contrast time-series were treated as evoked responses whose underlying sources were inferred per participant using the multiple sparse priors source reconstruction. The single-participant 3D source activity maps were entered into a group-level GLM to infer those sources which are differentially sensitive to the temporal expectations based on foreperiod and distribution. Thus, significant effects were inferred per time window using a paired F-test, contrasting the source estimates corresponding to the passage-of-time effect and the distribution effect, after thresholding the 3D (spatial) statistical parametric maps at *P* < 0.001 (peak-level, uncorrected) and correcting P-values for multiple comparisons based on cluster size at a family-wise error rate at *P* < 0.05. Sources were assigned anatomical labels using the SPM12 atlas provided by Neuromorphometrics, Inc.

### 2.7 Dynamic causal modelling

To explain the observed effects of temporal expectation related to foreperiod and tone onset distribution on the evoked neural responses in a biophysically realistic model, we used dynamic causal modelling (DCM) in a neural mass model based on a canonical microcircuit (Figure 3B; cf. Pinotsis, et al. 2013). Specifically, the observed effects were modelled as changes in effective connectivity between and within nodes of a network comprised of several cortical sources. In DCMs based on a canonical microcircuit architecture, the activity in each source is modelled using ordinary differential equations which describe the changes in postsynaptic voltage and current in four neural populations. These populations (spiny stellate cells in the granular layer, supragranular/superficial and infragranular/deep pyramidal cells, and inhibitory interneurons) are characterised by distinct profiles of ascending and descending connectivity both extrinsically (linking different sources) and intrinsically (coupling neural populations within a source). Specifically, there is a laminar asymmetry in the outputs of each source — for instance, superficial pyramidal cells propagate ascending signals to hierarchically higher areas, whereas deep pyramidal cells propagate descending signals to hierarchically lower areas. The output of each source is further modulated by intrinsic connectivity parameters, describing the strength of the self-inhibition of each neural population. The equations describing the dynamics at each source are provided below:

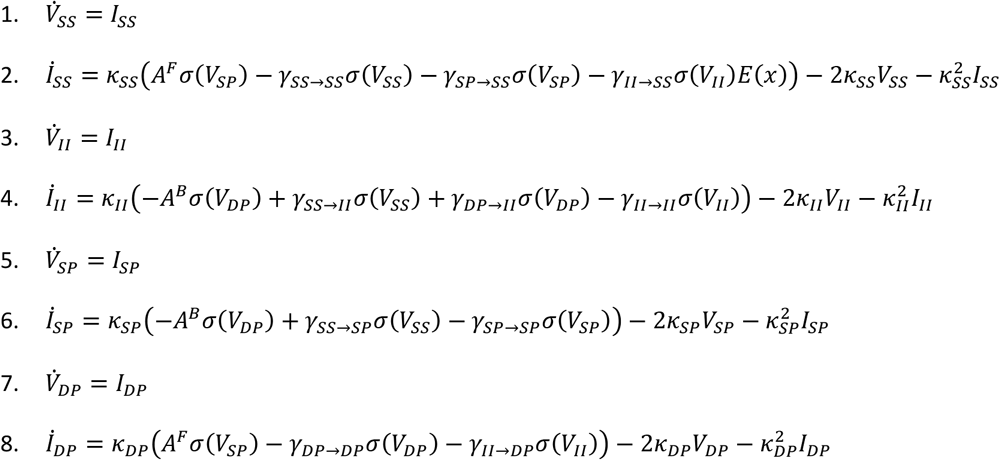

**Figure 3.**
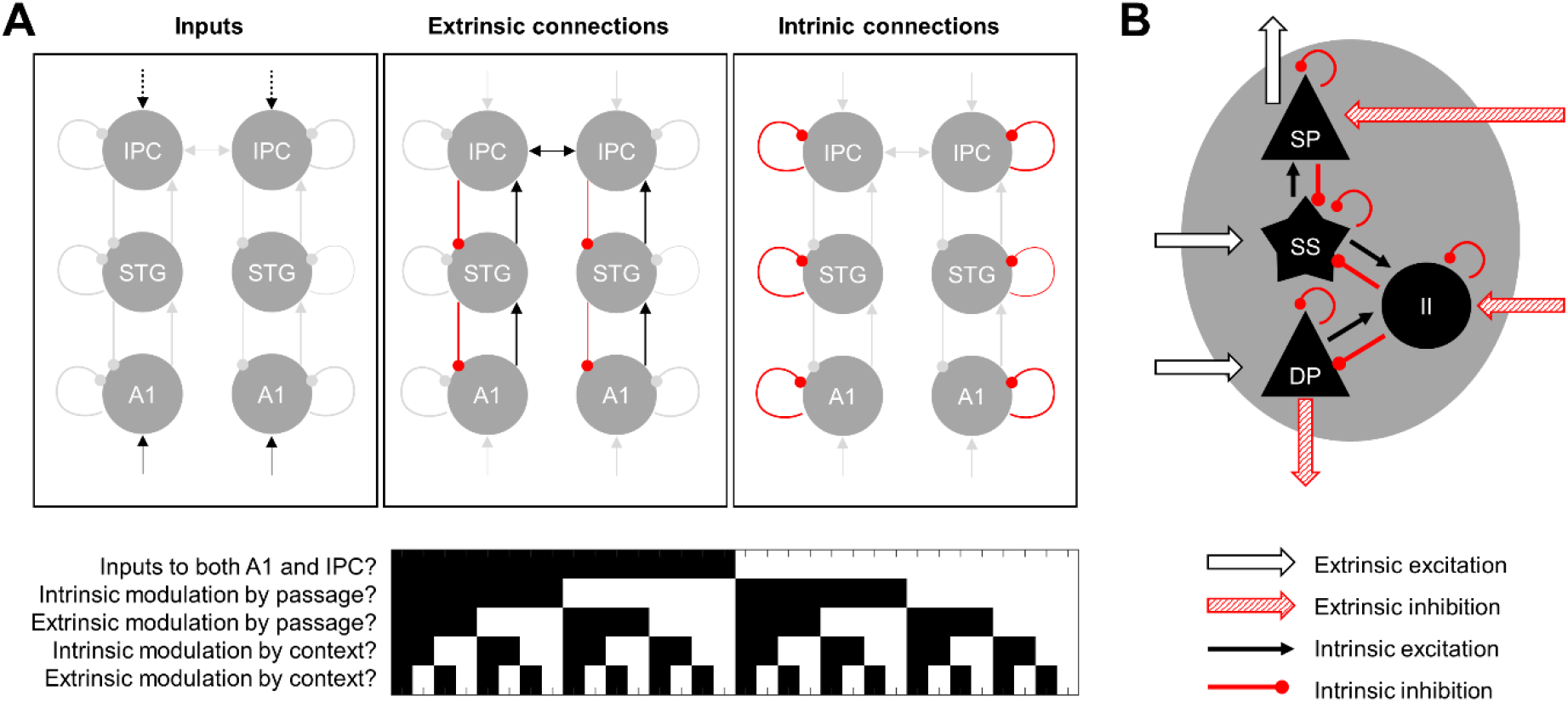
Dynamic causal modelling. ***(A)*** Model space: models had the same underlying architecture comprising 6 areas interconnected in an anatomically plausible way. The models differed with respects to which areas received external inputs (A1 only or A1 and IPC), and which connections (extrinsic vs. intrinsic) were modulated by each of the experimental manipulations (foreperiod vs. contextual expectation). Combinations of intrinsic modulations at the level of single regions are grouped for simplicity. This model space is depicted in the lower panel, with black/white shading representing absence/presence of a given input or modulation. ***(B)*** Each area corresponds to a microcircuit with 4 neural populations (SP: superficial pyramidal cells; DP: deep pyramidal cells; SS: spiny stellate cells; II: inhibitory interneurons) whose interactions determine the activity dynamics in the modelled network.

The distinct neural populations comprising a canonical microcircuit are indicated by subscripts SS (spiny stellate cells), II (inhibitory interneurons), SP (superficial pyramidal cells), and DP (deep pyramidal cells). The voltage and current of a population *m* are denoted by *V_m_* and *I_m_* respectively, determined by the synaptic rate constant *κ_m_s*. The sigmoid operator *C* transforms the postsynaptic potential into firing rate. The extrinsic ascending and descending connections are denoted by *A^F^* and *A^B^* respectively. Intrinsic connections from population *m* to *n* are denoted by *γ_m→n_*. Crucially, the intrinsic self-inhibition of superficial pyramidal cells *γ_SP→SP_* determines the gain of ascending connections, which has been linked to the precision of prediction error signalling in the predictive coding framework (Feldman and Friston, 2010). Finally, thalamic input *u* scaled by its weight *C* contributes to the changes in current of spiny stellate cells at the lowest level of the hierarchy. This model has been used in several other DCM studies of evoked responses (e.g., Moran, et al., 2013; Auksztulewicz & Friston, 2015; Brown & Friston, 2012), and neurophysiological inference using DCM has been validated in animal models (Moran, et al., 2011) and invasive recordings in humans (Papadopoulou, et al., 2015).

A model space containing two model families with 256 competing models each was designed to disambiguate alternative hypotheses regarding the connectivity patterns modulated by temporal expectation based on passage of time vs. distribution of stimulus events (Figure 3A). All models were based on the same basic network, comprising 3 bilateral sources identified in the source reconstruction of the observed effects (see Results 3.3), namely early auditory cortex (A1), superior temporal gyrus (STG), and inferior parietal cortex (IPC). The two families differed with respect to the subset of regions receiving driving (e.g., thalamic) input: in one family, the driving input was assumed to only reach bilateral A1 (cf. Auksztulewicz & Friston, 2015; Moran, et al., 2013), while in the other family the driving input reached both A1 and IPC, akin to non-sensory input regions for expectation signals modelled in other studies using DCM (Philips, et al., 2016; Gilbert & Moran, 2016). Each model allowed for a different subset of connections to be modulated by either of the experimental factors (passage of time and/or context). Specifically, we considered models in which each of the experimental factors could independently modulate the following subsets of connections: (1) A1 gain; (2) STG gain; (3) IPC gain; (4) extrinsic connectivity between regions. In the group-level analysis, first – for computational feasibility – we fitted only the full models (i.e., allowing for all connections to be modulated by both experimental factors) to each participant’s data. These models were fitted to the event-related fields in the 0-500 ms range relative to tone onset across all MEG channels, resulting in predicted sensor-level data (specifically, 8 principal spatiotemporal modes accounting for >99% of the variance across all sensors and time points). The reduced models (allowing for only a subset of connections to be modulated) were not fitted individually, but instead their parameter estimates (describing the effective extrinsic and intrinsic connectivity parameters) and model evidence (reflecting the model fit while penalising for model complexity) were obtained per participant using Bayesian model reduction of the respective full models (Friston, et al., 2016). Following this step, we obtained model evidence and parameter estimates for each model and participant. Second, to ensure that group-level model evidence or parameter estimates are not driven by outliers or models with a poor fit, we used parametric empirical Bayes to obtain group-level parameter estimates assuming that single-participant parameter estimates for each model come from a normal distribution (Friston, et al., 2016). Thus, we obtained group-level model evidence and parameter estimates for each model. Finally, to accommodate uncertainty inherent in comparing such a large number of models, we used Bayesian model averaging (Friston, et al., 2016) of the group-level models in the winning model family, which integrates single-model parameter estimates weighting them by the evidence of each respective model. The average model contained parameter estimates (mean and covariance estimates) used for statistical inference. Parameters were considered significantly modulated if they were different from 0 at a posterior probability threshold of 95% (Bonferroni-corrected across parameters).

## RESULTS

### 3.1 Behavioural results

Participants detected on average 95.6% of the deviants (SD = 5.3 across participants). For deviants presented in the second position, the waiting time (ISI) had a significant effect on reaction times (F_(2,46)_ = 9.29, p < .001; see Figure 1C), with deviant tones presented at shorter intervals after the first tone (ISI<500) being detected more slowly than deviants presented at the middle interval (ISI=500 ms; t_(23)_ = 3.33, p = 0.003; mean RT at short ISIs: 561 ms, medium ISIs: 499 ms) or at longer intervals (ISI>500, t_(23)_ = 4.97, p < .001; mean RT at long ISIs: 492 ms). We found no evidence for a difference between RTs to deviants presented at medium vs. long ISIs (t_(23)_ = 0.31, p > .05). However, given the relatively low number of trials with responses to deviants per condition (14 trials on average), the presence or absence of behavioural effects on the deviant tones should not be interpreted as an indicator for the presence or absence of neural effects on the standard tones (see below, 3.2).

### 3.2 Event-related fields – sensor space analysis

Both the foreperiod until the tone onsets and the distribution of tone onset times had an influence on sensor-level ERF amplitude. (Figure 4A). Specifically, the passage of time until tone onset (foreperiod) significantly modulated ERF amplitude between 267-500 ms with a bilateral topography. The distribution of tone onsets, on the other hand, had a significant effect on ERF amplitude later in time (between 427-500ms) and with a left-lateralised topography. We next wanted to know whether these clusters can be uniquely linked to foreperiod and temporal distribution expectations respectively, or, conversely, if they represent a mixture of both effects. To this end we repeated the analysis by (1) testing the effect of foreperiod while masking out the effect of onset distribution (masking threshold p<.001, uncorrected) and inferring the significant effects at p<.05, FWE-corrected; (2) testing the effect of onset distribution while masking out the effect of foreperiod (using identical thresholds); (3) testing the conjunction of foreperiod and distribution effects (thresholded at p<.001 and corrected for multiple comparisons at p_FWE_<.05). Here we found that the effects of foreperiod and context were independent of each other (Figure 4B). There was no evidence of overlap in the two clusters, as identified in the conjunction analysis (no conjunction cluster significant at a threshold of p<.001, after correcting for multiple comparisons at p_FWE_<.05).

**Figure 4.**
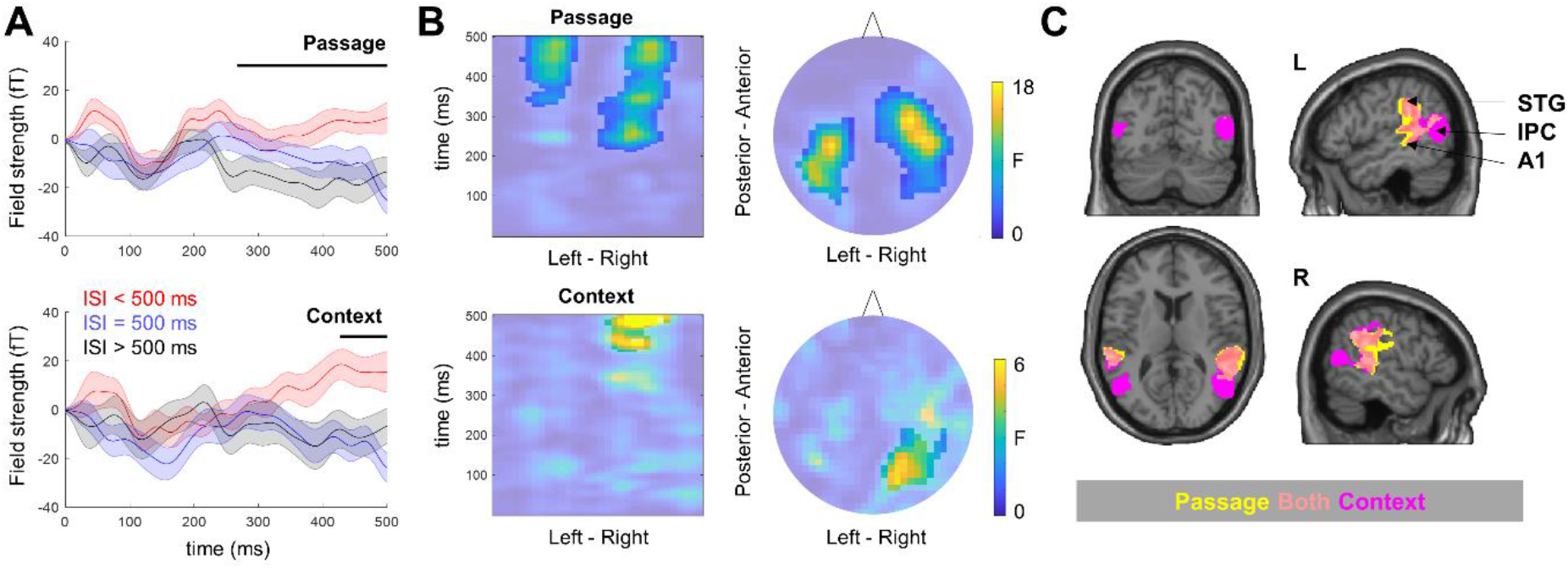
Differential effects of foreperiod and contextual expectation on evoked MEG responses. ***(A)*** Time-courses of averaged MEG amplitude extracted from channels showing significant effects of foreperiod (upper) and context (lower). Shaded areas denote SEM over participants. Black bar denotes time window of a significant cluster also shown in (B). ***(B)*** Both columns show maps of F statistics over parametric effects of passage (upper) and context (lower). Left column shows the time course of the effect on the Y axis. Right column shows the topography of the effect. The effect of foreperiod starts earlier than the effect of context. When thresholding at p<.001 and correcting using pFWE<.05, foreperiod has significant bilateral effects between 267-500ms while context has a left-lateralised effect between 427-500ms (unmasked areas). ***(C)*** Source reconstruction. In yellow, the significant (p<.001, pFWE<.05) difference between sources underlying passage and context in the long time window. In magenta, the significant (p<.001, pFWE<.05) difference between sources underlying passage and context in the short late time window. In orange, the overlap between the two. Slices centred at [±52 – 65 11] MNI coordinates. A1: primary auditory cortex; STG: superior temporal gyrus; IPC: inferior parietal cortex.

### 3.3 Dynamic causal modelling – source space analysis

Our next step was to reconstruct the sources of the sensor-level parametric effects of foreperiod and tone distribution expectation (Figure 4C; Table 1). This analysis revealed that for a long time window (250-500 ms), in which the effect of foreperiod modulated ERF amplitude, the power of source-level activity in bilateral early auditory cortices (A1) and superior temporal gyri (STG) significantly differed between temporal expectations stemming from foreperiod and tone onset distribution (voxels thresholded at p<.001 and corrected using pFWE<.05). Apart from these two bilateral regions, an analysis of the shorter, later time window (425-500 ms) in which ERF amplitude was modulated by expectation of tone distribution, revealed that bilateral sources including A1, STG, and additionally inferior parietal cortex (IPC) once again significantly differed between the two kinds of temporal expectation. In sum, IPC was uniquely sensitive to expectations of distribution of events in time, while A1 and STG were sensitive to both this distribution and the passage of time. We next used these 3 bilateral regions as a basis for subsequent dynamic causal modelling. The full models (with all connections allowed to be modulated by both kinds of temporal expectations) explained on average 50.64% (SD = 15.71%) variance of the individual-participant MEG channel x time x condition data. Figure 5B shows the average observed (measured) and predicted (modelled) data per channel and time point.

**Figure 5.**
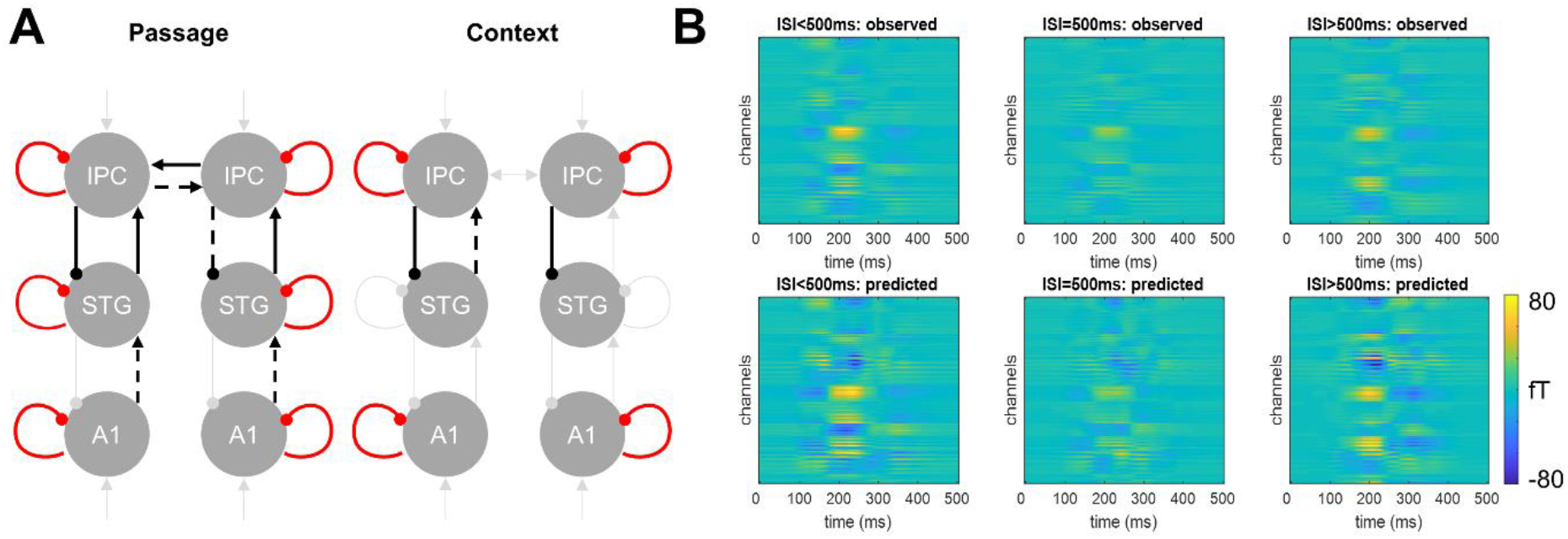
Dynamic causal modelling results. ***(A)*** Modulatory parameters of the winning model for the effects of passage of time (foreperiod; left) and context (right). Black: excitatory, Red: inhibitory, solid: stronger, dashed: weaker; light grey: connections not modulated by a given experimental factor. ***(B)*** Observed (upper row) and predicted (lower row) responses for each modelled condition (columns), time point (X axis), and MEG channel (Y axis).

**Table 1.**
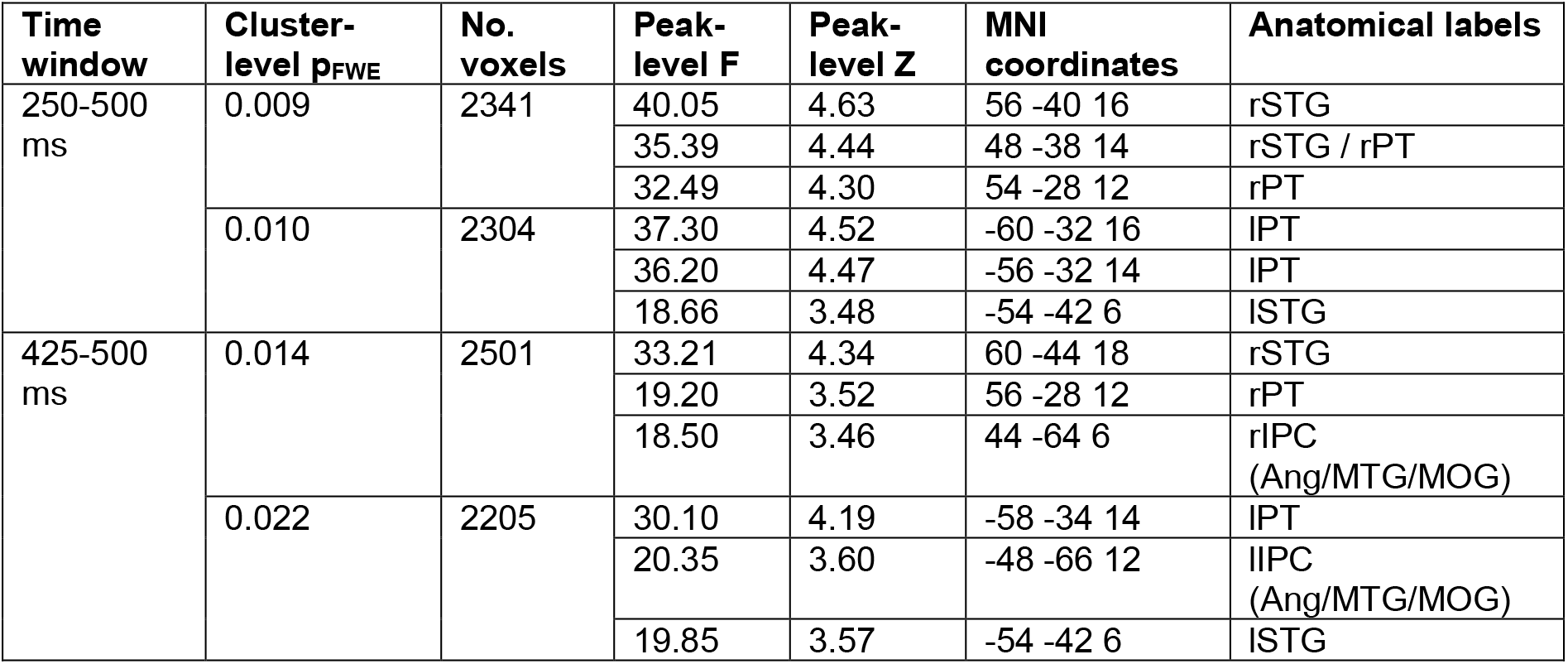
Source reconstruction results. STG: superior temporal gyrus; PT: planum temporale; IPC: inferior parietal cortex; Ang: angular gyrus; MTG: medial temporal gyrus; MOG: medial occipital gyrus.

Our model space included two families of models: in one family, inputs were assumed to only reach bilateral A1, representing ascending sensory information (cf. Auksztulewicz & Friston, 2015; Moran, et al., 2013), while in the other family the driving input reached both A1 and IPC. This second family was based on DCM literature suggesting that expectation signals are effectively modelled as inputs to non-sensory regions (Philips, et al., 2016; Gilbert & Moran, 2016), and indeed this family had higher model evidence on average in our dataset (posterior family probability >99.9%). In each family, the associated models differed with respect to the pattern of possible of extrinsic and intrinsic connections which would be modulated by foreperiod and/or temporal distribution. The Bayesian model average of all models in the winning family indicated that, while both passage-of-time and temporal distribution modulated the weight of extrinsic connections (linking different regions) and intrinsic connections (modelling gain within regions), their effects were dissociable at the level of specific connections both within and across regions. The posterior parameters of the winning model are plotted in Figure 5A and summarised in Table 2. These parameters indicate that, with stronger predictions of passage of time, the lower-order ascending (excitatory) connections between A1 and STG became weaker, while the higher-order ascending connections between STG and IPC became stronger. The descending (inhibitory) connections were modulated by passage of time only between higher-order regions (IPC and STG). The effects of passage of time on these descending connections were asymmetric, showing stronger inhibition in the left hemisphere and weaker inhibition in the right hemisphere. These effects were accompanied by an asymmetric modulation of lateral (cross-hemispheric) connections between left and right IPC, with net effective connectivity shifted towards the left hemisphere. The effects of expectation due to temporal distribution on extrinsic connections were limited to higher-order connectivity between STG and IPC, showing a left-lateralised decrease of ascending excitation and a bilateral increase in descending inhibition.

**Table 2.** DCM parameter estimates. Posterior means are reported on a logarithmic scale, e.g. a mean of −0.6631 corresponds to exp(−0.6631) = a decrease of the connection strength to 51.53% of the baseline due to a given experimental factor.

| Modulatory effect | Parameter | Posterior mean | Posterior probability | Description | Net effect |
| --- | --- | --- | --- | --- | --- |
| Foreperiod | B(1,1) | .3239 | >.9999 | A1 self-inhibition (left) | Inhibitory |
|  | B(3,1) | -.6631 | >.9999 | A1-STG connection (left) | Inhibitory |
|  | B(2,2) | .1038 | >.9999 | A1 self-inhibition (right) | Inhibitory |
|  | B(4,2) | -.2707 | >.9999 | A1-STG connection (right) | Inhibitory |
|  | B(3,3) | 0.2587 | >.9999 | STG self-inhibition (left) | Inhibitory |
|  | B(5,3) | 0.4014 | >.9999 | STG-IPC connection (left) | Excitatory |
|  | B(6,4) | 0.1655 | 0.9993 | STG-IPC connection (right) | Excitatory |
|  | B(3,5) | -.1621 | >.9999 | IPC-STG connection (left) | Disinhibitory |
|  | B(5,5) | .0960 | .9998 | IPC self-inhibition (left) | Inhibitory |
|  | B(6,5) | .2214 | .9999 | IPC cross-hemispheric (left-right) | Excitatory |
|  | B(4,6) | .2141 | >.9999 | IPC-STG connection (right) | Inhibitory |
|  | B(5,6) | -.3083 | >.9999 | IPC cross-hemispheric (right-left) | Inhibitory |
|  | B(6,6) | .0832 | .9996 | IPC self-inhibition (right) | Inhibitory |
| Context | B(1,1) | .1586 | >.9999 | A1 self-inhibition (left) | Inhibitory |
|  | B(2,2) | .6086 | >.9999 | A1 self-inhibition (right) | Inhibitory |
|  | B(6,4) | -.1775 | .9999 | STG-IPC connection (right) | Inhibitory |
|  | B(3,5) | .0888 | .9992 | IPC-STG connection (left) | Inhibitory |
|  | B(5,5) | .2643 | >.9999 | IPC self-inhibition (left) | Inhibitory |
|  | B(4,6) | .2718 | >.9999 | IPC-STG connection (right) | Inhibitory |
|  | B(6,6) | .3703 | >.9999 | IPC self-inhibition (right) | Inhibitory |

In addition to the dissociable effects on extrinsic connections, expectation due to foreperiod and temporal distribution also showed dissociable effects on intrinsic connections. Specifically, while increasing expectation due to foreperiod increased intrinsic connectivity in all regions bilaterally, expectation due to temporal distribution increased intrinsic connectivity in bilateral A1 and IPC but not in STG.

## DISCUSSION

Attending to events in time has been linked to a broad network of areas, reflecting the complexity of temporal expectation (Coull, et al. 2004; Muller and Nobre 2014). Here we investigated whether passage of time and tracking the distribution of events in time have dissociable neural substrates. We employed a two-tone paradigm where onsets of the second tone were normally distributed along five inter-stimulus intervals, and we looked at how the second tone in these pairs was processed as a function of elapsed time, and of the likelihood of the tone arriving at that particular inter-stimulus interval. We modelled passage of time and temporal distribution separately, with passage of time corresponding to a model where evoked activity reduced as the inter-stimulus intervals got longer (i.e. predictability grew), and distribution of events in time corresponding to a model where evoked activity was minimal at the middle interval (where predictability was greatest) but grew towards the edges of the distribution. We found that these two types of temporal expectation correlated with activity in a similar set of areas, but that the modulations began at different moments in time, and that they affected network connectivity in distinct ways.

Behaviourally, we found that tones that arrived before the most frequent, middle ISI, were paired with slower response times, confirming that likelihood had an effect on how tones were processed. A recent study with normally distributed tone onsets also found faster responses to tones with more predictive foreperiods (Herbst, Fiedler, and Obleser 2018), which echoes previous findings (Hsu, Hämäläinen, and Waszak 2013; Coull and Nobre 1998; Miniussi, et al. 1999). On the neural level, since the tones were presented quickly one after the other (with the ISIs ranging from 250 ms to 750 ms), we first applied convolution modelling to estimate the evoked field to the second tone while controlling for the influence of the first tone. After this step, we were able to include early, middle and late arriving tones into a single search for cortical sources. We found two clusters of activity, one beginning early (starting at 267 ms after stimulus onset) and corresponding to passage of time, and one beginning late (starting at 427 ms after stimulus onset) and corresponding to temporal distribution of tone events. Importantly, we found that the neural effects related to these two types of temporal expectation were separable from each other.

We then localised the effects of temporal expectation to sources corresponding, bilaterally, to the primary auditory cortex (A1), superior temporal gyrus (STG) and inferior parietal cortex (IPC). While A1 and STG were sensitive to both this distribution and the passage of time (corresponding to the early and late clusters of activity in the ERF time-series analysis), IPC was uniquely sensitive to expectations of distribution of events in time (corresponding specifically to the late activity clusters). In previous research, A1 has been implicated not only as a hub for auditory processing, but also as a modality-independent timekeeper (Kanai, et al. 2011). The STG and IPC have, similarly, often been found to be modulated by temporal expectation (Morillon and Baillet 2017; Wiener, Turkeltaub, and Coslett 2010; Rao, Mayer, and Harrington 2001; Cui, et al. 2009; Coull, Nazarian, and Vidal 2008). One area that is also frequently found to be involved in time estimation, is the supplementary motor area (SMA) (Herbst, Fiedler, and Obleser 2018; Bueti, et al., 2010; Coull, et al., 2008; Akkal, et al., 2004). It has even been suggested that processing of time has motor origins (Morillon and Baillet 2017). Interestingly, we did not observe SMA activity to be involved in our temporal hazard. Potentially this might be because the temporal processing in our study was implicit: although it aided task performance, this was a pitch discrimination task, and not one that involved estimating durations. However, one other study with a tone discrimination task involving similarly distributed tone onset times also found SMA activity (Herbst, Fiedler, and Obleser 2018). In that study, SMA was involved in foreperiod tracking in blocks with flat (i.e. equiprobable) onset time distributions but not in blocks with strong contextual expectations of the distribution. It is therefore likely that our search for cortical sources, which was specifically sensitive to areas involved both in passage of time and contextual distribution of events in time, led to SMA being excluded.

We next looked at how each of these two types of temporal expectation modulated connectivity within and between A1, STG and IPC. We found that both types of temporal expectation affected the strength of connections between regions in the auditory network. We were particularly interested in testing whether changes in connection weights would be in line with predictive coding, namely, corresponding to decreases in feedforward excitation, increases in feedback inhibition and increases in intrinsic inhibition when a tone is more predictable. However, the changes in connectivity profiles that were affected by temporal expectation were more varied, especially when it came to tracking predictability as a function of elapsed time.

The most prominent common feature between the two types of temporal expectation was how they modulated intrinsic connections in bilateral A1, STG and IPC. With growing tone onset probability, whether due to passage of time or due to their distribution, intrinsic connections consistently led to greater inhibition of neural activity (i.e. lower gain). Gain modulation has indeed been posited as a core feature of predictive processing (Garrido, et al., 2009; Auksztulewicz & Friston, 2016). In the context of repetition suppression, when an initially novel (unpredicted) stimulus starts repeating and forming a standard (predicted) stimulus, neural gain – modelled as intrinsic connections – shows a relative decrease following the first repetitions (when a prediction is being established) and a gradual rebound with later repetitions (when a prediction is fully formed; Garrido, et al., 2009). A similar change in intrinsic connectivity was evident here in the entire network for predictability related to elapsed time, and in A1 and IPC, but not STG, for predictability associated with temporal distribution. A number of previous studies have indicated a role for the STG in temporal processing. For instance, BOLD activity evoked by target onset in both STG and SMA has been shown to reflect the cumulative hazard of a target appearing at a given moment, independent of modality (auditory vs. visual) or the presence of a motor response (Cui, et al., 2009). Importantly, when participants were informed of the exact target onset in that study, BOLD signal no longer correlated with the foreperiod, suggesting that activity in these regions did not simply reflect passage of time, but rather a temporal probability estimate. Another study found that STG (but not A1 or IPC) was involved in estimating relative stimulus duration (Coull, et al., 2008). Therefore, a plausible explanation for the difference in STG gain modulation between the two types of temporal expectation is that a more involved time estimation was necessary for estimating the expected peaks of temporal distribution in our study, which required activity in STG, preventing self-inhibition.

The added complexity of assessing temporal distribution (relative to passage of time) was also visible in the fact that predictability due to distribution was largely tracked in higher-level connections between IPC and STG. Differences in prediction complexity have previously been linked to hierarchical differences in prediction and prediction error signalling, with hierarchically higher areas integrating information over longer time scales and representing more complex predictions (Auksztulewicz & Friston, 2016; Chao, et al., 2018; Kiebel, et al. 2008). In the current study, fully in line with predictive coding, tones that were more predictable based on the distribution (i.e. middle ISI tones relative to both early and late tones) led to less feedforward activation from STG to IPC, and more feedback inhibition from IPC to STG.

Similarly, if the prediction came from monitoring elapsed time, less excitation was fed forward from A1 to STG. The same study that found reduced gain during prediction formation also found a monotonically decreasing feedforward excitation from A1 to STG, as well as a reduction in ERPs, in repeated standard tones (Garrido, et al. 2009). Although there was no manipulation of stimulus timing, the similarity with our experiment lies in stimulus predictability: standard tones are predictable and expected, especially when repeated multiple times, and therefore less forward excitation likely mediates reduced prediction error signalling at lower levels of the cortical hierarchy (Auksztulewicz & Friston, 2016).

However, at higher levels of the cortical hierarchy, the changes in connectivity begin to diverge from what predictive coding models would posit. Firstly, more (not less) excitation was fed forward from bilateral STG to IPC after the occurrence of a more predictable tone. Secondly, when it comes to feeding inhibition back from IPC to STG, passage of time led to an opposite effect in the two hemispheres, with inhibition increasing with stronger predictions in the left hemisphere, but decreasing in the right hemisphere, amounting to stronger net activation in the right IPC and weaker activation in the left hemisphere. Thirdly, stronger predictability with passage of time led to an asymmetric modulation of cross-hemispheric connectivity between the left and right IPC, amounting to the right IPC exerting stronger influence on the left IPC than vice versa. While stimulus expectation is most often linked to a reduction in forward signalling, expectation enhancement has also occasionally been observed (Segaert et al., 2013; Recasens et al., 2014) and the underlying mechanisms are still under debate (Auksztulewicz & Friston, 2015). For instance, the net effect of expectation on neural activity may be mediated by non-specific response dampening or by selective response sharpening (de Lange, et al., 2018); furthermore, expectation of different features may be mediated by qualitatively different cortical circuits (Auksztulewicz, et al., 2017). Interestingly, the reversing of prediction-related inhibition of neural activity by other factors such as attention has been described before (Kok, et al., 2012). In the current study, lower-level inhibition of ascending signalling may reflect a non-specific dampening of prediction error signalling as passage of time increases the predictability of a received stimulus; higher-level ascending excitation, on the other hand, may reflect a signal halting the ongoing time estimation as the stimulus has been received and can be processed by other regions, including the attentional network. A right-hemispheric IPC asymmetry has previously been reported specifically in a paradigm based on encoding time intervals (Rao, Mayer, and Harrington 2001) and linked to the lateralisation of the frontoparietal attentional network.

The differences between how two types of temporal expectation modulated connectivity suggest that, while stronger predictions *can* lead to less feedforward excitation and more feedback inhibition, these changes are not passed along the entire cortical hierarchy. In fact, they are more likely to be contained between a smaller number of cortical areas that are directly specific to a certain type of information processing, with, in this case, the more complex predictability related to tone onset distribution, modulating only higher order connections but not influenced by changes in activation coming from the primary auditory cortex. In addition, at the higher levels of the hierarchy the prediction errors were not suppressed by prediction related to passage of time, but instead changed the connectivity pattern in a different, often opposite way. Taken together, our results show demonstrate that different aspects of temporal expectation can be dissociable in terms of the latency of neural responses, the underlying sources and connectivity patterns, leading to a more nuanced view of how prediction and prediction error signalling may be expressed in cortical circuits.

## ACKNOWLEDGMENTS

We would like to thank Kia Nobre and Floris de Lange for their helpful comments on experimental design and data analysis. R.A. is funded by the European Commission’s Marie Skłodowska-Curie Global Fellowship (750459).

